# Combining Multi-Site/Multi-Study MRI Data: Linked-ICA Denoising for Removing Scanner and Site Variability from Multimodal MRI Data

**DOI:** 10.1101/337576

**Authors:** Huanjie Li, Stephen M. Smith, Staci Gruber, Scott E. Lukas, Marisa M. Silveri, Kevin P. Hill, William D. S Killgore, Lisa D. Nickerson

## Abstract

Large multi-site studies that pool magnetic resonance imaging (MRI) data across research sites or studies, or that utilize shared data from imaging repositories, present exceptional opportunities to advance neuroscience and enhance reproducibility of neuroimaging research. However, both scanner and site variability are confounds that hinder pooling data collected across different sites or across different operating systems on the same scanner, even when all acquisition protocols are harmonized. These confounds degrade statistical analyses and can lead to spurious findings. Unfortunately, methods to address this problem are scant. In this study, we propose a novel denoising approach for multi-site multimodal MRI data that implements a data-driven linked independent component analysis (LICA) to efficiently identify scanner/site-related effects for removal. Removing these effects results in denoised data that can then be combined across sites/studies to improve modality-specific statistical processing. We use data from six different studies collected on the same scanner across major hardware (gradient and head coil) and software upgrades to demonstrate our LICA-based denoising approach. The proposed method is superior compared to the existing methods we tested and has great potential for large-scale multi-site studies to produce combined data free from study/site confounds.

## Introduction

Neuroimaging studies aimed at understanding healthy brain structure and function as well as diseases of brain structure and function have proliferated with the ubiquitous availability of magnetic resonance imaging (MRI) as a tool for neuroscience research. While the vast majority of neuroimaging studies have been conducted within a single research site, it is now being recognized that pooling multi-site data, either through large-scale single and multi-site trials or through data sharing initiatives, has great potential for obviating some of the shortcomings of single-site studies. For example, increasing sample size by pooling multi-site MRI data can greatly enhance the statistical power to detect subtle effects or to study rare conditions and can enable the analysis of subgroups within a cohort (Button et al., 2013; Varoquaux, 2017). Pooling data also offers increased reliability and confidence about the size of an effect by averaging out unforeseen biases of individual studies and taking advantage of the wider variety of patient types and disease etiologies that are represented in multi-site studies (Van Horn et al., 2009). Neuroimaging data sharing is of increasing importance in the field of neuroimaging. As described in a Special Issue of NeuroImage titled “Sharing the wealth: Neuroimaging data repositories”, as of 2015, there were more than 40 imaging data sharing repositories, with repositories having specific datasets ranging from very small sample sizes to very large sample sizes. Eickhoff et al. (2016) present a table of repositories that are described in more detail throughout the special issue. Pooling the shared data from multi-site would present exceptional opportunities to advance neuroscience and enhance reproducibility of neuroimaging research. However, methodological issues related to combining multi-site/study data are not addressed in by any of the brief reports included the Special Issue.

Currently, there are more than a dozen ongoing NIH-funded large-scale neuroimaging studies, including the Adolescent Brain Cognitive Development (ABCD) Study (https://abcdstudy.org), a long-term study of brain development with 21 study sites collecting data in a target of approximately 10,000 children that just released the first wave of data in 4,500 participants. The NIH has also funded the disease connectomes (on anxiety, depression, psychosis, and other health conditions, http://www.humanconnectome.org/disease-studies), which are partially harmonized with the Human Connectome Project (HCP). The HCP study collected 4 hours of multimodal MRI data in more than 1,100 healthy young adults (at a single site). These connectomes present the opportunity for combining data across those studies. In the United Kingdom, the UK Biobank is conducting the world’s largest health imaging study with the aim of improving the prevention, diagnosis and treatment of a wide range of serious and life-threatening illnesses. Imaging data will be collected in 100,000 participants in the coming years, with imaging being conducted across three sites (http://www.ukbiobank.ac.uk/2016/09/brain-imaging-results-from-5000-subjects/). The first wave of UK Biobank data in 5,000 participants has also been released (Miller et al., 2016).

Neuroimaging data from these all of these initiatives will ultimately be made available to other researchers via data repositories. While the strength of these large-scale studies and of existing imaging data repositories lies in combining multi-site data to create large datasets that overcome limitations of small neuroimaging studies, combining images obtained from different sites/studies presents formidable challenges. Inconsistent MRI data collection platforms, imaging parameters, or other study differences may introduce systematic variability that can confound the true effect of interest and make the interpretation of results obtained from combined data difficult (Czanner et al., 2006; Jovicich et al., 2005). This is not only a problem for combining multi-site or multi-study data, but also for long-term longitudinal studies, including those with data collection at a single research site. In this case, the scanner may undergo hardware or software upgrades, which are by design intended to improve the performance of the scanner and could thus result in differences in signal to noise or contrast to noise over a prolonged study period, an effect that is compounded in multi-site longitudinal studies (Takao et al., 2011; Venkatraman et al., 2015). Glover et al. (2012) provide an in-depth discussion of how various factors such as field strength and scan parameters can lead to unwanted inter-site variability. They also provide recommendations for the design and execution of multi-site MRI studies, including numerous recommendations for the statistical analysis of such data, thus combining data across sites must be conducted with great care to ensure that sources of between-site variation are minimized and that the presence of residual between-site variation is taken into account in higher level statistical modeling.

Many studies have investigated the effects of scanner/site-related variability in cross-sectional and longitudinal data and found clear differences in high spatial resolution structural imaging (Focke et al., 2011; Isca et al., 2015; Keihaninejad et al., 2010; Littimann et al., 2006; Takao et al., 2011), diffusion tensor imaging (DTI) (Bartsch et al., 2014; Huisman et al., 2006; Kochunov et al., 2015; Pagani et al., 2010; Vollmar et al., 2010) and functional MRI (fMRI) (Casey et al., 1998; Zivadinov and Cox, 2008; Costafreda et al., 2007; Friedman et al., 2008; Wegner et al., 2008; Zou et al., 2005) outcomes across scanners. These studies reported scanner effects resulting from a number of factors, including field strength, scanner vendors, and major scanner upgrade. For example, Takao et al. (2011) evaluated the effects of an upgrade in scanner hardware (referred to as drift) and inter-scanner variability on global and regional brain volume changes using longitudinal data obtained on two scanners and found that scanner drift and scanner variability obscured actual longitudinal brain volume changes as evidenced by comparing the observed longitudinal brain volume changes to findings from previous studies of aging and brain volume. Using a small sample of subjects who had each completed scan sessions at four different study sites, Jovicich et al. (2009) demonstrated that multi-site MRI data collected using scanners from different vendors and with different field strengths introduced a bias in mean volume differences.

For DTI data, several studies have shown that DTI measurements (i.e., fractional anisotropy (FA) and mean diffusivity (MD)) are highly site-specific and that scanner manufacturer differences influence the reliability and quantification of DTI measurements (Pohl et al 2016; Venkatraman et al., 2015). Multi-site fMRI studies have also shown that group differences in brain activation during tasks and functional connectivity of resting-state networks might be attributable to scanner hardware differences, software differences and multi-site variations (Casey et al., 1998; Zivadinov and Cox, 2008). On the other hand, some studies have found that scanner effects did not interfere with the effect of interest. For example, Stonnington et al. (2008) found that different scanners and upgrades had negligible effects on segmented gray matter images in a multi-site study of Alzheimer’s disease. Han et al. (2006) showed that a scanner upgrade neither increased variability nor introduced bias in automated measurements of cortical thickness. Furthermore, Jovicich et al. (2009) showed that a scanner upgrade did not significantly change the variance between structural MRI data collected pre- and post-upgrade, although the upgrade may have introduced bias in the estimated mean volume differences. For example, for a Siemens Trio TIM upgrade, there was a significant reduction in the variance of the reproducibility of the measured volumes of some structures, including caudate, putamen, and pallidum among others, post-upgrade. However, the sample size was small in both the Han et al. (2006) and Jovicich et al. (2009) studies, therefore, these results may not be extrapolated to other pulse sequences and scanners not specifically investigated in these two reports.

Approaches to minimize the effects of multiple scanners/sites when combining MRI data have been proposed for each MRI modality, i.e., structural MRI data (Chen et al 2014; Fennema-Notestine et al., 2007; Keihaninejad et al., 2010; Pardoe et al., 2015; Takao et al., 2011), DTI data (Pohl et al 2016; Venkatraman er al., 2015; Fortin et al., 2017) and fMRI data (Feis et al., 2015; Glover et al., 2012; Griffanti et al., 2014; Salimi-Khorshidi et al., 2014). For statistical analyses, two techniques have emerged as the most common approaches to combining MRI data across studies/sites. The first, and perhaps most common approach, is the inclusion of site/study covariates in the higher-level general linear model (GLM) implemented to assess group differences (Glover et al., 2012; Chen et al., 2014; Fennema-Notestine et al., 2007; Venkatraman et al., 2015). For example, Fennema-Notestine et al. (2017) suggested that linear regression analyses with inter-site effects modeled as a random effect should be applied to remove multi-site effects when pooling multi-site structural MRI data. However, Glover et al. (2012) recommend treating inter-site effects as random effects only when the number of sites is large (i.e., six or more). For fewer sites, they recommend treating these effects as fixed effects. Venkatraman et al. (2015) used a linear mixed-effects model to estimate the differences in measurements of FA and MD between scanners both within and across field strengths (i.e., one single 1.5T and three different 3.0T Philips scanners) in a training dataset of healthy older adults; they then applied a GLM with the scanner differences from the training data as covariates to correct for inter-scanner effects in a new cohort of healthy subjects. The found that reliability across field strengths for FA and MD were low, with mean intra-class correlations of 0.35 and 0.31 in white matter, respectively. Using linear correction factors greatly increased reliability across scanners for most regions, but did not fully correct the systematic differences between scanners.

A second approach that has been used to a lesser extent is modality-specific independent component analysis (ICA) for denoising. ICA is a data-driven technique that attempts to decompose a multivariate signal into independent non-Gaussian signals (Bell and Sejnowski, 1995), and identify spatiotemporal patterns that reflect signals of interest versus processes that are related to motion and other artifacts. Chen et al. (2014) applied ICA to multi-site structural MRI data and investigated the associations of the resulting spatiotemporal patterns with various scanning parameters to assess the influence of site on the data. Components that were related to scanner effects were then eliminated from the original data to “denoise” the data. They also compared the ICA-based denoising approach with the standard approach of including site as a covariate in the GLM and found that both the GLM approach and the ICA-based denosing method mitigated scanner effects, however, the ICA-based approach was better able to handle collinear effects.

The over-arching goal of these methods for addressing scanner/site effects from imaging data is to first develop an accurate “model” of the effects and then to remove or statistically control for those effects, typically through regression. In the case of the GLM approach, the “model” is a simple regressor comprised of values corresponding to site/scanner/study in the group-level model to statistically control for potential differences *post hoc,* and in the case of ICA-based denoising, the “model” is the spatial pattern of a noise-related independent component. Inherent in both methods is that denoising (or statistically controlling for) scanner/site variability is implemented based on individual modalities (i.e., structural, DTI and fMRI). Furthermore, because of the many technological advances in hardware and scanning sequences, it is now possible to collect data for many different modalities during a single imaging session. Structural, diffusion, task and/or resting state fMRI data are now routinely collected in participants within 30-60 minute scanning sessions and the inclusion of multimodal sequences may create new problems when combining multi-site/study data, as each modality may be differentially impacted by scanner/site effects, and therefore could require different corrections.

In the present study, we propose a novel multimodal MRI denoising method based on data fusion of multiple MRI measurements to identify multimodal spatial patterns related to scanner and/or study variations that can be used to denoise these effects from each individual modality. In the first step of our proposed approach, linked ICA (LICA) (Groves et al., 2011, 2012) is applied to, for example, morphometric maps, FA, MD and MO (diffusion tensor outcomes), and brain activation or connectivity maps all together to decompose the data into spatial maps and corresponding subject loadings that capture sources of variability in the data. While LICA (and other data fusion methods) have so far only been applied to investigate neurobiology, we develop LICA as a tool for denoising site/study sources of variance to facilitate combining MRI data. We capitalize on the strengths of LICA, namely that by using all measurements for every subject together in the LICA, we can better identify scanner/site effects as patterns that are distinct from meaningful patterns of brain structure/function than doing ICA on each modality separately. For example, the LICA more efficiently models the common variance among the multiple measurements made in each subject, which improves its ability to identify between-subject sources of variability in the data and to separate out interesting between-subject effects from uninteresting effects such as those related to site/study variability. Once scanner/site effects have been identified, they can be removed from the original measurements to provide a “clean” set of measurements for each modality that are free of study/site effects that can be used for further statistical analyses.

In the present study, we demonstrate how LICA can be applied to remove study-related sources of variance from multimodal MRI data collected across six different studies from a single site. As expected, the LICA identified multimodal components associated with uninteresting scanner/study effects denoised using three different regression-based denoising methods for removing scanner/study-related components from the original measurements. The LICA components are a set of multimodal MRI spatial patterns and a corresponding set of subject loadings, which are measures of the strength of the multimodal spatial pattern present in each individual. Crucially, the subject loadings were used to identify scanner/site-related noise components by testing the associations of the loadings for each component with scanner/site variables. Once the noise components were identified, either the spatial patterns or the loadings for these noise components were used for denoising. In the first denoising method, a single multivariate regression was implemented to regress the subject loadings for all noise components against the subject series for each modality (e.g., the 4D file constructed by concatenating all subject’s spatial maps for a given modality into a single file) to remove the noise effects (LICA-R1). The second method applied dual regression (Filippini et al., 2009; Nickerson et al., 2017) to remove noise components by regressing the LICA spatial maps of all components against each modality’s subject series to obtain subject-specific regression weights for all components; subject-specific regression weights of noise components were then regressed against the subject series to remove the noise effects (LICA-R2). LICA-R1 and R2 are “hard” aggressive denoising methods that remove any shared variance between the signal and noise components when these two are not completely orthogonal. In addition to the hard aggressive methods, we also tested a third “soft” regression-based denoising method using the subject loadings for all components, LICA-SR, which regressed out only the unique variance associated with the noise components (Griffanti et al., 2014).

The data we used to test our proposed method contained similar sources of variability as might be observed in multi-site/study data as they were collected using the same scanner, but with different acquisition parameters, software versions, and spanned a major hardware upgrade of a Siemens Trio to a TIM Trio, which may introduce differences analogous to using different scanner models.

## Methods

### Study Data

Data from 179 subjects collected from six different studies were used for the present work. Initial quality assurance to assess for problems with the structural images (which are required for good registration of all modalities to standard space), head motion, artifacts, and other problems in any modality resulted in the removal of data from 23 subjects. Thus, MRI data from 156 subjects were used for the present study. Data included high-resolution structural images, diffusion data, and fMRI data, although not all data were collected in each participant, as each study collected data from different modalities. This resulted in “missing data” for some subjects for some modalities when considering all modalities together. Data from 73 chronic heavy marijuana (MJ) users (near daily use and positive THC screen on test day) (age = 24.6 ± 6.8, 62 male and 11 female) and 83 non-MJ using healthy controls (HC) (age = 24.2 ± 5.2, 50 male and 33 female) were included in our analyses.

All data were collected using the same Siemens 3T Trio, but with 3 different scanner software versions (SSWV) and over the course of a major hardware upgrade. Versions VA23A and VA25A were used prior to a major hardware and software upgrade of the Trio (TIM upgrade), while VB17A was used post-TIM. Acquisition sequences also differed across the studies, thus, these data contain similar sources of variability as multi-site/multi-study data, including scanner (with pre-post TIM essentially representing two different Siemens scanners), software versions, and study protocol.

### MRI acquisition parameters

High-resolution structural MRI data for all six studies were collected using a T1-weighted MPRAGE with 128 slices and flip angle (FA) = 12°. Studies 1 and 5 utilized echo time (TE)/repetition time (TR) = 2.74 ms/2100 ms, 1.5 × 1.0 × 1.3 mm^3^ voxel size. Studies 2, 3 and 4 utilized TE/TR = 2.15 ms/2000 ms, 1.5 × 1.0 × 1.3 mm^3^ voxel size. Study 6 utilized TE/TR = 2.25 ms/2100 ms, 1.0 × 1.0 × 1.0 mm^3^ voxel size.

Diffusion MRI data for studies 2 and 3 were collected using an echo-planar imaging (EPI) sequence with scan acquisition parameters: TE/TR = 89 ms/9300 ms, 60 slices, 2.0 × 2.0 × 2.0 mm^3^, 48 directions, b-value = 700 s/mm^2^ (three b = 0 images without diffusion weighting were acquired).

Task fMRI data for studies 2, 3, 4 and 6 were collected with gradient-echo EPI. Acquisition imaging sequences for studies 2, 3 and 4 were: TE/TR/FA = 30 ms/3000 ms/90°, 40 coronal slices, 3.1 × 3.1 × 5.0 (no gap) mm^3^ voxel size, interleaved acquisition. For study 6, TE/TR/FA = 30 ms/2000 ms/90°, 34 coronal slices, 3.5 × 3.5 × 3.5 (no gap) mm^3^ voxel size, with interleaved acquisition. fMRI data were collected while participants performed a multi-source interference task (MSIT) of inhibitory processing during the fMRI scan. The MSIT task is described in more detail in Gruber et al. (2012).

### Data processing

All data processing was done using FMRIB Software Library, FSL (Smith et al., 2004 https://fsl.fmrib.ox.ac.uk/fsl/fslwiki/), FreeSurfer (https://surfer.nmr.mgh.harvard.edu) and Matlab (Mathworks, Inc., Natick, MA). Optimized modality-specific preprocessing pipelines, including quality assurance to identify data with excessive motion or other artifacts, were used to produce standard-space outcome images for each subject for a given modality, including: 1) modulated grey matter (GM) images generated by FSL-VBM (Douaud et al., 2007), 2) vertex-wise cortical thickness (CT) and pial surface area (PSA) maps estimated using FreeSurfer by means of automated surface reconstruction scheme (Dale et al., 1999; Fischl and Dale, 2000; Fischl et al., 1999a, 1999b, 2001, 2002, 2004), 3) FA, MD and tensor mode (MO) images calculated using FSL FDT (Smith et al., 2006), and 4) MSIT brain activation maps estimated by FSL FEAT (https://fsl.fmrib.ox.ac.uk/fsl/fslwiki/FEAT) analysis of the MSIT task fMRI data. For GM and fMRI data, the images were registered to 2 × 2 × 2 mm^3^ MNI standard space; FA, MD and MO images were registered to 1 × 1 × 1 mm^3^ MNI standard space. For each outcome that was derived from the structural MRI, DTI and MSIT fMRI data, a “subject series” was created by concatenating the resulting spatial maps across all participants into a single 4D (volume x subjects) data file. These outcomes are hereafter also referred to as modalities even though they are derived quantities.

### Structural MRI

GM images: For all high-resolution structural MR images that passed through quality assurance (e.g., with no significant motion or artifacts observed in the structural images), non-brain tissue was removed, followed by gray matter-segmentation and registration to the MNI 152 standard space using non-linear registration via FNIRT (Anderson et al., 2007a,b). The resulting images were then averaged together and flipped along the x-axis to create a left-right symmetric, study-specific gray matter template. All native gray matter images were then non-linearly registered to this study-specific template and “modulated” to correct for local expansion (or contraction) due to the non-linear component of the spatial transformation. Modulated GM images were then smoothed using an isotropic Gaussian kernel with a sigma of 4 mm.

CT and PSA images: Using the same high-resolution structural MR images that were used for GM, CT measurements were obtained by reconstructing representations of the gray/white boundary and the pial surface and then calculating the distance between the surfaces at each vertex across the cortical mantle (Dale and Sereno, 1993; Dale et al., 1999). PSA was estimated by registering each subject’s reconstructed surfaces to a common template, and the relative amount of expansion or compression at each vertex was used as a proxy for regional arealization. PSA were resampled and mapped to a common coordinate system using a non-rigid high-dimensional spherical averaging method to align cortical folding patterns (Fischl et al., 1999a, 1999b, 2008). Both CT and PSA images were smoothed with a Gaussian kernel (full width of half maximum = 10 mm).

### DTI data

FSL’s FDT and TBSS tools were used to generate FA, MD, and MO maps for white matter for each subject (Smith et al., 2006). FSL eddy was used to correct DTI data for motion and eddy currents (Andersson and Sotiropoulos, 2016). After removal of non-brain tissue, fitting of diffusion tensors on corrected data was applied using *dtifit* to compute FA, MD, and MO maps (Smith, 2002). Each subject’s FA map was transformed into common space by warping to the FMRIB58_FA template using FMRIB’s nonlinear image registration tool (FNIRT, Andersson et al., 2007a,b). The mean FA volume over all individuals was computed and then thinned to create a mean FA skeleton. The mean skeleton was thresholded at FA exceeding 0.2 to minimize partial volume effects at the boundaries between tissue classes. Individual FA maps were multiplied by the mean skeleton to compute skeletonized FA maps for each subject. MD and MO maps for each subject were processed similarly to result in a standard space skeletonized FA, MD, and MO map for each subject.

### MSIT fMRI data

The fMRI data was processed to estimate subject-specific task spatial maps (TSM) corresponding to brain activation during the interference condition relative to the control condition using a first-level GLM implemented in FSL FEAT v6.00. Each fMRI dataset underwent quality assurance to identify excess subject motion and other artifacts followed by standard pre-processing, which included: motion correction, slice-timing correction, non-brain tissue removal, spatial smoothing using a Gaussian kernel of FWHM 7.0 mm, grand-mean intensity normalization of the entire dataset and high-pass temporal filtering with a cut-off of sigma = 42.0 s. Registration of fMRI data to high-resolution structural images was carried out using FLIRT (Jenkinson et al., 2002). Registration of high-resolution structural images to standard space was first done using FLIRT, then further refined using FNIRT nonlinear registration. Time-series statistical analysis was carried out via GLM with regressors corresponding to the inhibitory control condition convolved with a gamma hemodynamic response function using FILM with local autocorrelation correction (Woolrich et al., 2001). The Z-stat image from the first-level analysis (the contrast of parameter estimate map for interference – control) was thresholded at Z = 2.3 and -2.3, resulting in the subject-specific TSM. These images were registered to standard space using the FLIRT/FNIRT transformation matrices in a single step.

### LICA-based Noise Components

Subject series were constructed for each imaging outcome, or modality, by concatenating images for that modality across subjects, using the same subject order for each modality. To account for the fact that participants from different studies had different measurements, a volume of zeros was used to represent a subject’s “missing” data for a given modality. This is a simple way to fix the subjects’ noise precision to zero for that modality in the LICA algorithm, which gives the missing data for that modality no weight in the ICA decomposition (Groves et al., 2012). Thus, seven 4D datasets, one for each of the seven modalities, were constructed with 156 subject volumes for each modality. In applying LICA, participants with high noise variance (1/precision) will influence a significant fraction of the LICA components (Groves et al., 2012). We identified several patterns that were driven by individual subjects in the first-pass of the LICA even though the LICA algorithm is supposed to automatically down weight these subjects in determining the updates for the LICA component (Groves et al., 2012). Since these participants (N=23) drove several components, and were thus outliers, we removed them from the analysis and re-ran LICA. This is similar in practice to handling outliers when conducting group independent component analysis (GICA) of fMRI data. Namely, GICA maps can reveal problems in the subject data that lead to corruption of GICA maps – these data must be remediated or removed to obtain improved GICA maps. All data from these 23 participants (who all only had structural MRI outcomes) were removed, resulting in data from 133 subjects being included in our final analysis (62 chronic heavy MJ users and 71 non-MJ using HC). Table 1 shows details of the 133 subject “dataset” that was used, including MRI scan-types that were collected for each of the six studies, the number of subjects from each study, and the SSWV information.

**Table 1.** For each study, the number of HC and MJ subjects with each modality is shown (upper and lower row for each modality for each study, respectively). Also shown is the total number of HC/MJ and all subjects for each modality, with SSWV in parenthesis (VA23A/VA25A/VB17A). --indicates that data for that modality were not collected for that group (HC/MJ) in the study. HC: healthy control; MJ: marijuana; SSWV: scanner software version. VA23A and VA25A are pre-TIM software versions, VB17A is post-TIM software version. Missing data indicates how many participants do not have data for that modality. Publications from the original studies provide additional details about each study and the findings from the original data.

| Study | Structural | DTI | MSIT fMRI | Publication |
| --- | --- | --- | --- | --- |
|  | HC (VA23A/VA25A/VB17A )<br>MJ (VA23A/VA25A/VB17A) |  |  |  |
| 1 | 7(4/3/--)<br>8(5/3/--) | --<br>-- | --<br>-- |  |
| 2 | 16(--/9/7)<br>21(--/10/11) | 15(--/8/7)<br>20(--/9/11) | 16(--/9/7)<br>15(--/8/7) | Gruber et al., 2012,<br>2014, 2015 |
| 3 | --<br>16(--/--/16) | --<br>16(--/--/16) | --<br>14(--/--/14) | Gruber et al., 2012,<br>2014, 2015 |
| 4 | --<br>9 (--/--/9) | --<br>-- | --<br>9 (--/--/9) | Hill et al., 2017 |
| 5 | 13(--/13/--)<br>8(--/8/--) | --<br>-- | --<br>-- | Mashhoon et al.,<br>2015 |
| 6 | 35 (--/--/35)<br>-- | --<br>-- | 30 (--/--/30)<br>-- | Cui et al., 2015 |
| Total HC | 71 (4/25/42) | 15 (--/8/7) | 46 (--/9/37) |  |
| Total MJ | 62 (5/21/36) | 36 (--/9/27) | 38(--/8/30) |  |
| Total All | 133<br>(9/46/78) | 51 (--/17/34) | 84 (--/17/67) |  |
| Missing Data | 0 | 82 | 49 |  |

The subject-series for all 7 modalities were analyzed simultaneously using LICA implemented in Matlab (https://fsl.fmrib.ox.ac.uk/fsl/fslwiki/FLICA). LICA identifies spatial components, with each component being an aggregate of spatial patterns from each modality, along with a set of subject loadings, one for each component. Loadings for a given component are shared between all of the modalities represented in that component, and indicate the degree to which that multimodal component is present in any individual subject. The subject loadings for each component were assessed for relationships with SSWV, study, and participant variables (demographic, drug use, and task performance measures) using linear regression. Those components whose subject loadings related *only* to SSWV/study variability were identified as noise components. We ran the LICA decomposition at several different dimensionalities (L) of 12, 14, 15, 17, 20, 25 and 30 components to “tune” the LICA to identify the dimensionality that resulted in components that were stable across most dimensionalities and that had loadings most clearly associated with SSWV/study (and not with subject variables). Resulting multimodal spatial patterns from LICA are converted to pseudo-Z-statistics by accounting for the scaling of the variables and the SNR for individual modalities. LICA spatial maps can then be thresholded at Z = 2.3 for visualization, which is based on assuming an explicit spherical noise model during the LICA decomposition (Groves et al., 2012). Notably, LICA can identify both multimodal components and modality specific components (comprised of effects from only a single or small number of modalities), which makes it well-suited to identifying scanner/site/study effects that may be common to all modalities or that may uniquely affect only some modalities.

### Data Denoising

Once the LICA components associated with scanner/study effects are identified, they can be used to denoise the original data. We used multivariate regression for data denoising. Similar to Griffanti et al. (2014), who tested “hard” regression (which removes noise effects fully from the data by removing all shared variance that may exist between signal and noise) and “soft” regression (which removes only the unique variance associated with noise) for denoising fMRI data, we tested three approaches we developed for denoising the multimodal MRI data using the LICA results:

### LICA-R1 denoising: Hard regression using LICA subject loadings

A single multivariate regression of the subject loadings for only the noise components against the original data was implemented to remove noise components from each modality as follows:

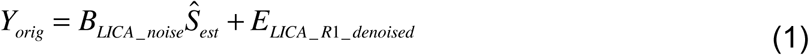

*Y*_*orig*_ is the 4D subject-series for a given modality reorganized into a 2D data matrix (M subjects x N voxels), *B*_*LICA_noise*_ are the matrix of subject loadings for the noise components identified from the LICA (M subjects x P components). 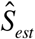 are the estimated spatial maps of regression coefficients (P components x M subjects). *E*_*LICA_R1_denoised*_ are the residuals from the regression, which represent clean data using the LICA-R1 de-noising method. Each modality was denoised separately.

**LICA-R2 denoising: Hard regression via dual regression using LICA spatial maps** A multivariate dual regression procedure (Nickerson et al., 2017) using the LICA component spatial maps was implemented to remove noise components as follows:

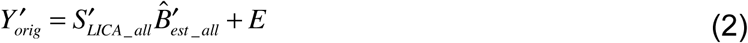

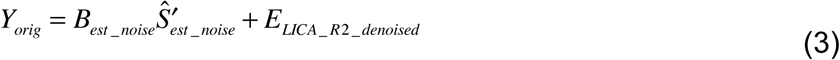

*Y*_*orig*_ is the original 4D data, reorganized into a 2D data matrix, similar to above, with the prime indicating transpose. In Eq. 2, *S*_*LICA_all*_ are the spatial maps of all the components from the LICA. 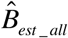 are the estimated regression coefficients for all the components based on the spatial map. *B*_*est* _ *noise*_are the estimated regression coefficients for noise components, which are similar but not equal to the subject loadings for the noise components. In the second regression (Eq. 3), *B*_*est* _ *noise*_from 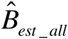 is regressed against *Y*_*orig*_.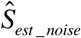 are the estimated spatial maps of regression coefficients. *E*_*LICA_R2_denoised*_are the residuals from the regression, which represent clean data using the LICA-R2 denoising method. Each modality was denoised separately.

### LICA-SR denoising: Soft regression using LICA subject loadings

First, multiple regression of the full set of LICA subject loadings against the original subject-series was done to estimate the contribution of all components (i.e., signal and noise). The unique variance of the noise was removed by subtracting the contribution of the noise components from the original data (Griffanti et al., 2014):

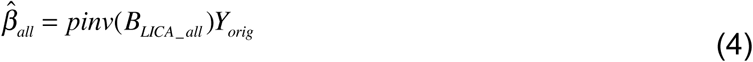

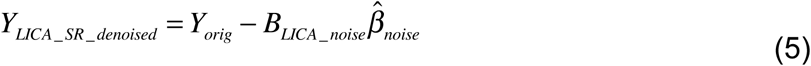

*B*_*LICA_all*_ are the subject loadings of all the components from the LICA. 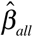 are the estimated contributions of both signal and noise components. *B*_*LICA_noise*_ are the subjectn loadings of the noise components and 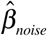 are the regression coefficients. *Y*_*LICA_SR_denoised*_ represents the clean data using LICA-SR. Each modality was denoised separately. All regressions in Equations 1-5 were easily implemented using FSL command line tools (fsl_glm and fsl_regfilt).

### Other methods for combining multi-site/study data

We compared the performance of LICA-R1/R2/SR for denoising our multi-study data with two other approaches that have been proposed to address scanner confounds when combining MRI data across studies/sites: a higher-level GLM with a site/study covariate included in the group-level model, and modality-specific ICA-based denoising.

For group-level GLM regression, the parameters corresponding to SSWV and study were included as the regressors in the design matrix. The scanner and study variables were coded as categorical variables, similar to coding used in ANOVA factor effects models. For example, the scanner variable has three levels (i.e., VA23A, VA25A and VB17A), thus we use two coding variables to represent the scanner variable. VA23A was chosen as the reference level, thus for each coding variable, rows of the design corresponding to VA23A will have -1. The first coding variable is -1 for VA23A, 1 for VA25A and 0 for VB17A; the second coding variable is -1 for VA23A, 0 for VA25A and 1 for VB17A. Voxel-wise regression of scanner and study effects against the original data was done using the fsl_glm command line tool.

ICA-based denoising of individual modalities was done using FSL MELODIC (Multivariate Exploratory Linear Optimised Decomposition of Independent Components – Beckmann and Smith, 2004) with automatic dimensionally estimation. Noise components were identified based on the associations of extracted independent component “time series”, which in this case are subject loadings, with SSWV and study variables. Once all noise components were identified, fsl_regfilt was used to remove the noise components.

### Assessment of the performance of LICA denoising methods

We demonstrate how data denoising works for individual modalities by constructing artificial datasets based on splitting existing data into groups based on, e.g., pre- and post-TIM upgrade or study, then assessing group differences pre- and post-denoising. Since the groups are artificially constructed by scanner/study variables, any group differences may be attributed to scanner/study and denoising should result in minimizing these effects and a subsequent reduction of these group differences (to zero ideally). For structural MRI and fMRI modalities, healthy control data were used to construct two groups of healthy subjects based on either pre-/post-TIM or by study parameters. GM and CT maps from healthy controls were separated in two groups based on SSWV (pre- vs. post-TIM. For pre-TIM, age = 24.0 ± 5.2 years, 10 females and 19 males; for post-TIM, age = 25.2 ± 5.5 years, 20 females and 22 males). As almost all the TSM were from data collected post-TIM, TSM maps from healthy controls were divided in two groups based on different acquisition parameters: Group 1 comprised data from studies 2, 3, and 4, which used the same acquisition parameters (age = 23.3 ± 3.6 years, 10 females and 6 males), and Group 2 comprised data from study 6 (age = 25.1 ± 4.9 years, 16 females and 14 males), which used different acquisition parameters from the studies in Group 1. For DTI outcomes, there were only 15 HC subjects, thus we were not able to construct groups to assess denoising of DTI measures.

Group differences in each modality were assessed for statistical significance before and after denoising using two-group *t*-tests with threshold-free cluster enhancement (TFCE, Smith and Nichols, 2009) in FSL’s Randomise (https://fsl.fmrib.ox.ac.uk/fsl/fslwiki/Randomise) with 5,000 permutations to achieve a significance level of *p* = 0.05, corrected for family-wise error. Regions that showed statistically significant group differences were selected as regions of interest (ROI) to illustrate associations between group differences and SSWV and study variables in these areas before and after denoising.

## Results

From the final LICA, three noise components had loadings that were strongly associated with SSWV and/or study variables, and not related to any other variables, (Figs. 1-3). While LICA dimensionality L = 15 identified these three noise components with loadings that were most clearly associated with SSWV/study, the spatial patterns were also observed at different dimensionalities (e.g., the LICA spatial patterns were stable). A component that was heavily weighted by FA and MD (Fig. 1) had loadings that were strongly associated with both SSWV (*p* = 1.91e-08) and study (*p* = 1.88e-04). This component showed widespread effects in FA and MD and region-specific effects in GM, TSM, CT, and PSA. Each modality had different contributions to this component, i.e., the weights for each modality were: FA (29%), MD (61%) MO (3%), GM (1%), TSM (0%), CT (5%) and PSA (1%).

**Figure 1.**
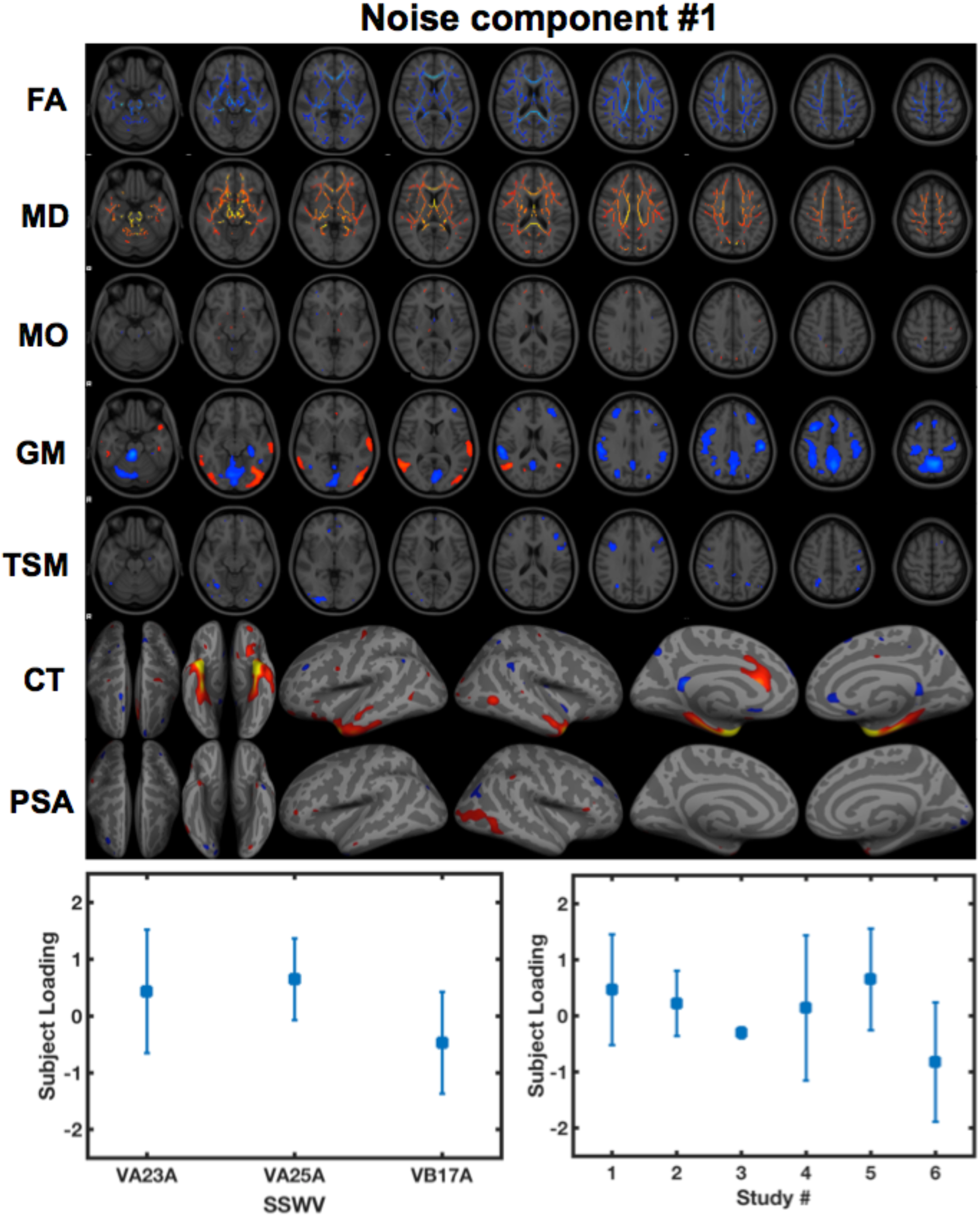
A multimodal noise component identified from LICA. Subject loadings for this component were strongly associated with both SSWV (*p* =1.91*10^-8^) and STUDY (*p* = 1.88*10^-4^) variables. The spatial pattern shows global effects in FA and MD and region-specific effects in GM, fMRI, CT and PSA. The spatial maps were thresholded at Z = 2.3.

**Figure 2.**
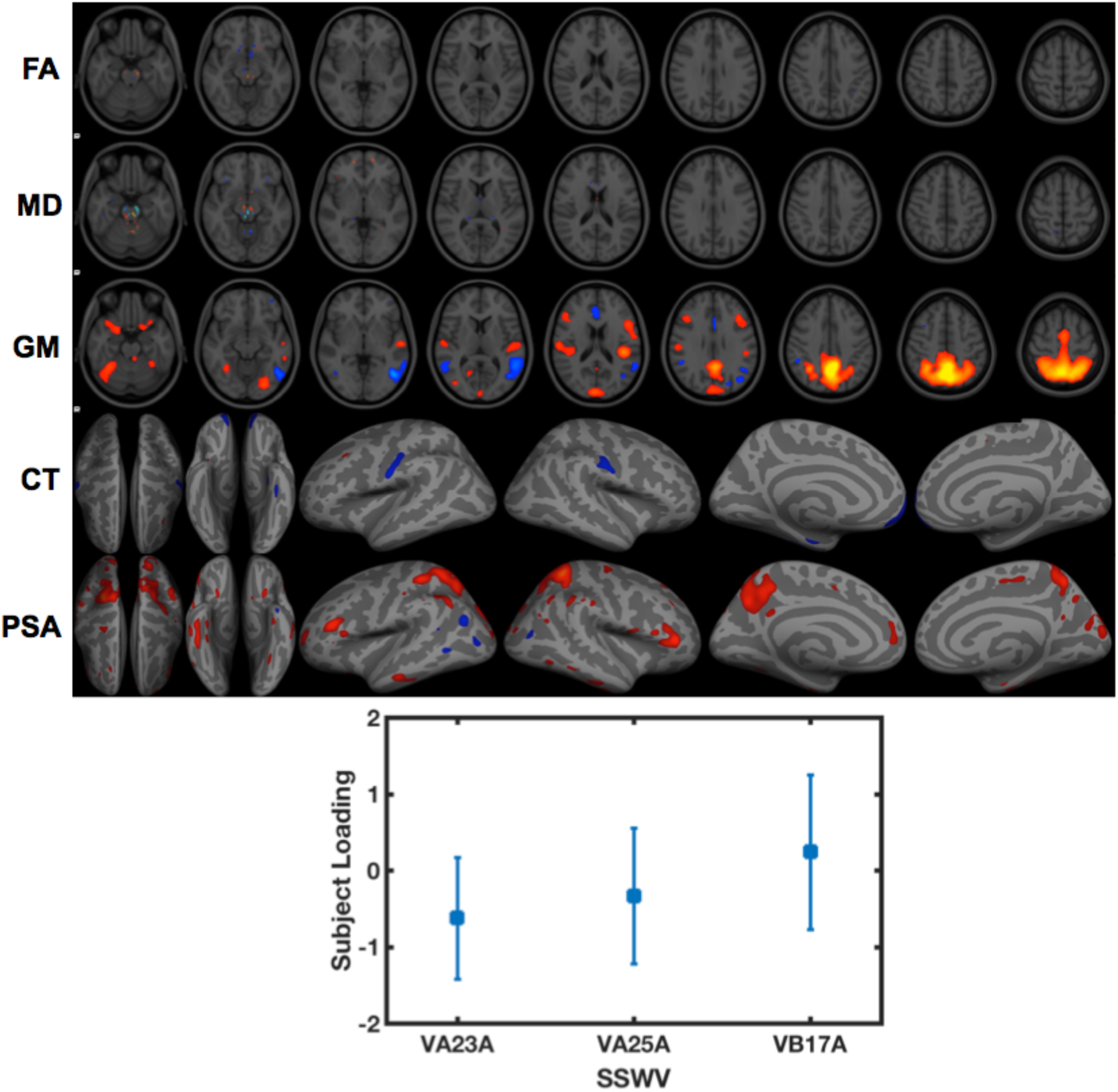
A second multimodal noise component identified from LICA. Loadings for this component were only associated with SSWV (*p* = 2.15*10^-4^). The spatial pattern shows region-specific effects in FA, MD, GM, CT and PSA. The spatial maps were thresholded at Z = 2.3.

**Figure 3.**
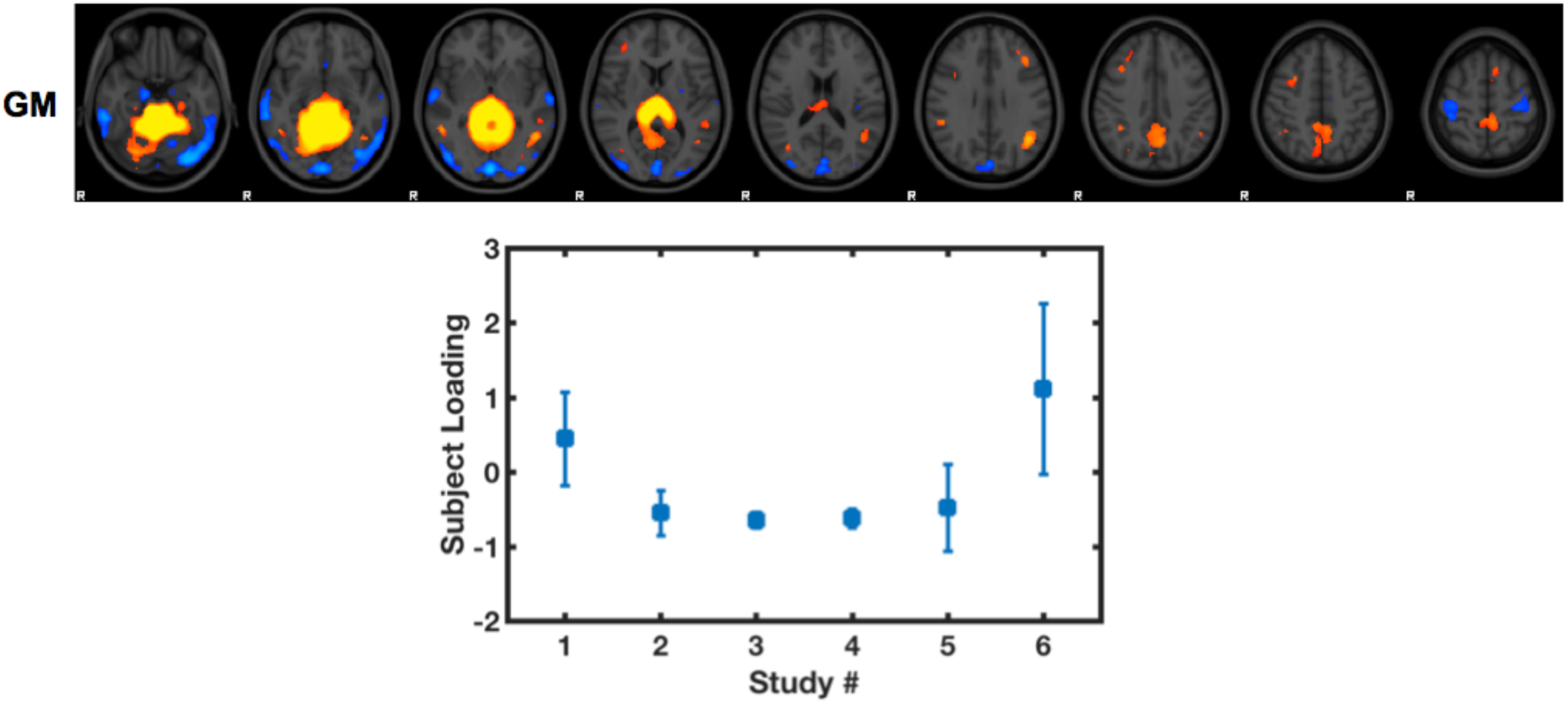
A third noise component identified from LICA. Loadings for this component were only associated with STUDY (*p* = 1.17*10^-6^). The spatial pattern was weighted very high for GM, with no or small weights for the other modalities (which are not shown). The spatial map was thresholded at Z = 2.3.

A second multimodal component was identified that had a strong linear association (*p* = 2.15e-04) with SSWV. This component showed region-specific effects in FA, MD, GM, CT and PSA (Fig. 2), with weights for each modality given by FA (3%), MD (11%), GM (31%), TSM (2%), CT (12%) and PSA (36%), indicating a more “multimodal” effect of SSWV. A third component heavily weighted to GM only (Fig. 3) was identified, with loadings that were strongly related to study (*p* = 1.17e-06).

Fig. 4 shows the results of the two-group *t*-test to assess differences in GM between the two HC groups, before and after denoising. Non-denoised results were obtained using the original GM data without any noise variable regression or data denoising. Therefore, the first row in the figure corresponds to the group differences in GM associated with pre- and post-TIM upgrade that would be obtained using a standard higher-level GLM. Group differences were observed in widespread brain regions, including insula, sub-cortical brain regions, and occipital cortex. Many of these regions are also seen in the GM spatial maps from the noise components identified using LICA, especially in Fig. 3 (the values in the LICA maps are reversed, however, they are in the same direction when considering the loadings). While it is not known exactly what these differences are due to, statistically controlling for SSWV and study using GLM regression (shown in the second row) is modestly effective at removing SSWV effects, with some remaining noise-related differences and the appearance of new regions with spurious group differences.

**Figure 4.**
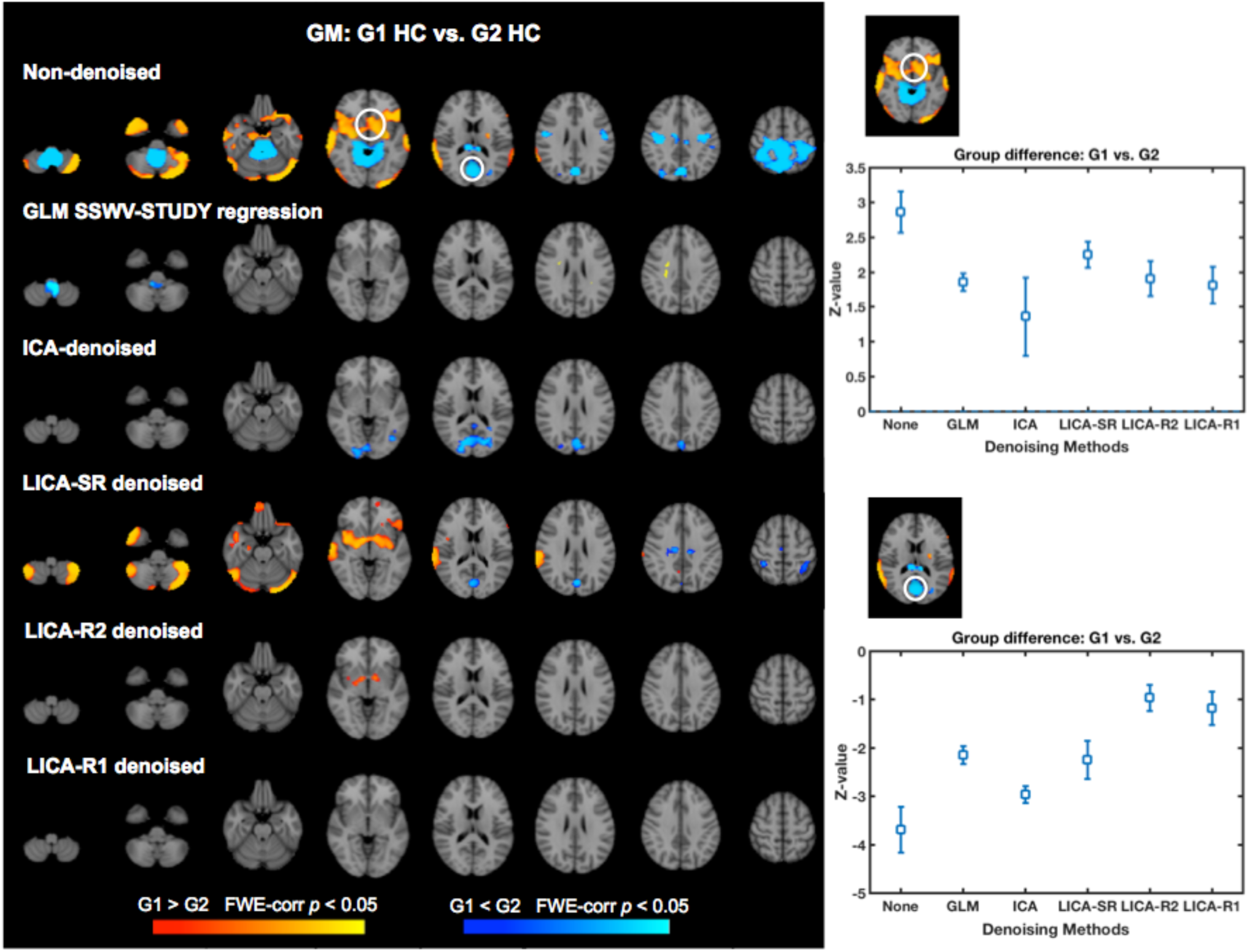
Group-level analysis of GM maps before and after data denoising. Data were constructed for two HC groups, each with different SSWV. G1 contains 26 subjects with data acquired pre-TIM upgrade; G2 contains 42 subjects with data acquired post-TIM upgrade. The first row shows the results of the group comparison without any data de-noising or regression (i.e., using original GM data). The second row shows statistically significant group differences with SSWV-STUDY regression. The third row shows group differences after GM maps have undergone modality-specific ICA denoising. The bottom three rows show the group differences for GM data that have been denoised with LICA-SR, -R2, and -R1 methods. All spatial maps are thresholded at *p* < 0.05, corrected. The plots on the right show the mean Z-value and standard deviation from voxels with significant group differences shown in the white circles (first row). Group differences (Z-values) are greatest with no denoising and are reduced in both regions (e.g., toward Z = 0) using any of the denoising methods, although LICA-based methods show a clean spatial map, suggesting this method has the best performance.

The third row shows the results using modality-specific ICA-denoising. A total of 34 components were identified using automatic dimensionally estimation, and among them, twenty components were found to have subject loadings that were significantly related to SSWV and study variables. Thus, these 20 components were removed from the original data using regression (Equations 1-5). Rows 4-6 show the results using the LICA-SR, LICA-R2 and LICA-R1 denoising methods to remove the three noise components from LICA that had loadings that were strongly associated with SSWV and study variables (shown in Figs. 1-3). Modality-specific ICA-denoising, LICA-SR and LICA-R2 denoising methods were also modestly effective at denoising, removing most of the effects that were related to SSWV and study variables with some remaining noise-related differences still apparent. The hard aggressive denoising methods LICA-R2/R1 show better denoising performance than the soft regression-based denoising method LICA-SR. Strikingly, the LICA-R1 denoising method reduced all SSWV- and study-related effects, while not introducing false positives. Thus, LICA-R1 denoising demonstrated superior performance for denoising scanner/study effects from GM data. Among the regions significantly associated with SSWV, we randomly chose two of them to illustrate the denoising performance of each method. The mean Z-value and the standard deviation of voxels within the ROI were calculated (from the outputs of the two-group *t*-test) to illustrate the effectiveness of each method for removing SSWV effects (e.g., to show that apparent differences in the maps are not simply due to thresholding effects). In this case, Z-values for non-denoised data reflect the differences due to SSWV/STUDY, and the Z-values for the denoising methods should move closer to 0 if the denoising method is working well. The LICA-based methods show more effective performance over both ROIs, with LICA-R1 perhaps having the best as assessed subjectively by looking at the whole map. E.g., the mean Z-value plots suggest that the LICA-based methods are similar in performance, even though LICA-R1 is the only method resulting in no significant differences in the spatial map.

Fig. 5 shows the group differences in TSM between the two HC groups before and after data denoising. The procedure for each denoising method was the same as for the GM analyses. For modality-specific ICA, 48 components were estimated using automatic dimensionally estimation, and among them, 10 components were found to be significantly related with SSWV and study variables suggesting that SSWV/study may have less effect on this modality (which is also consistent with the LICA components – which do not show strong effects for fMRI TSM and with the first row of Fig. 5, which shows only a few areas that are affected). Thus, these 10 components were removed. The denoising performance of each method for fMRI outcomes was consistent with the performance of each method for GM. With no denoising, there were regions with statistically significant group differences that were statistically significantly associated with study and SSWV variables. GLM regression removed noise related group differences from many regions. ICA, LICA-SR and LICA-R2 denoising methods remove most SSWV/study effects and reduced group differences in most regions. LICA-R1 appears to perform slightly better than other methods with all noise-related group differences no longer significant after denoising.

**Figure 5.**
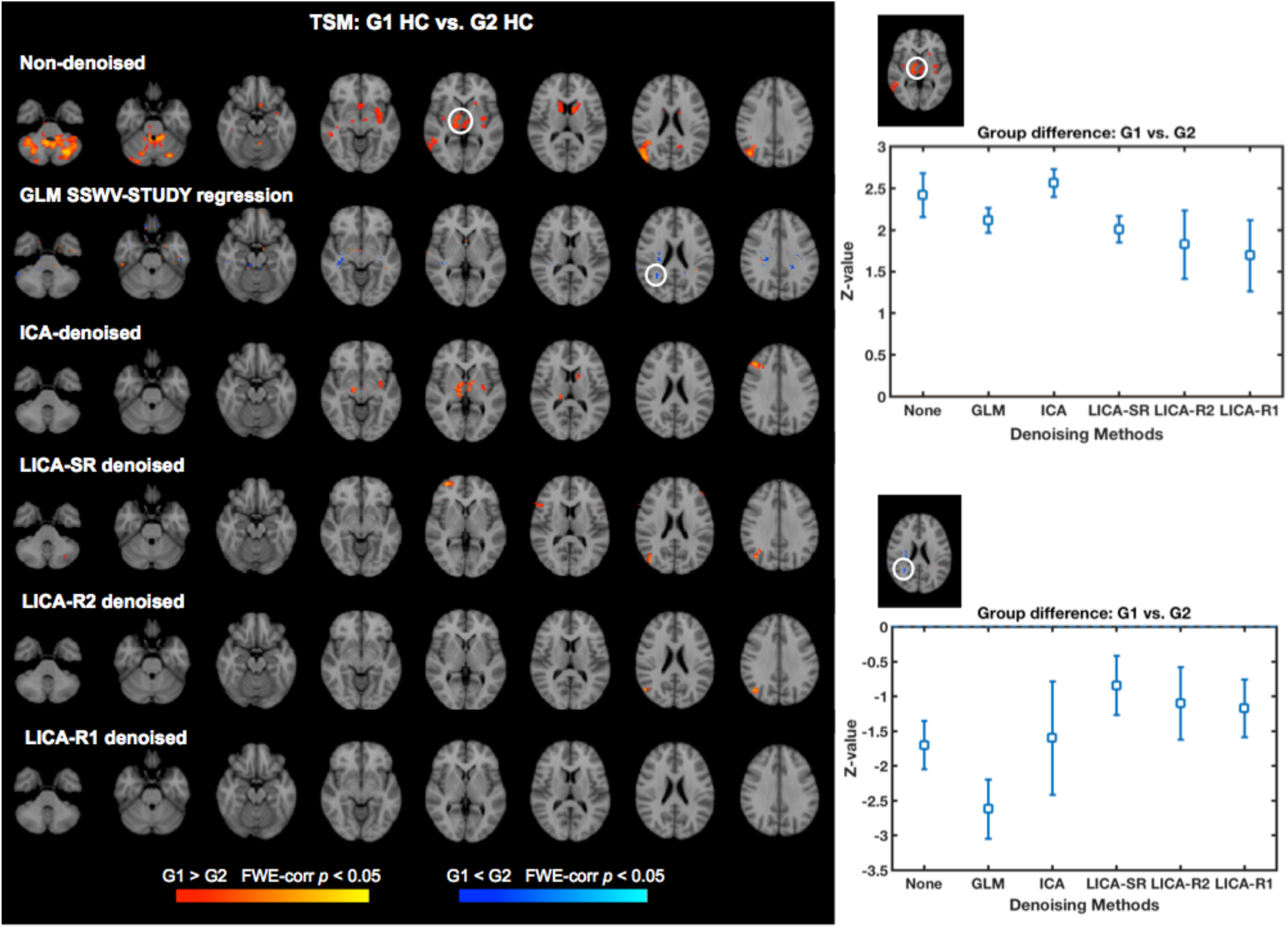
Group-level analysis of fMRI task spatial maps (TSM) before and after data denoising. The two HC groups were constructed based on SSWV. G1 contains 16 subjects with data acquired pre-TIM upgrade; G2 contains 30 subjects with data acquired post-TIM upgrade. Resulting group difference maps without any data denoising (first row) and with data denoising (rows 2-6) and plots (right) of the mean Z-values and standard deviations extracted from voxels with significant group differences (shown in the white circles) show that LICA-R1 denoising of TSM achieves the best noise removal.

Fig. 6 shows the group differences in CT between the two HC groups before and after data denoising. The procedure for each denoising method was the same as for GM/TSM analyses. The denoising results are also quite consistent with the GM and fMRI denoising results. For ICA-denoising, only one component was detected by automatic dimensionally estimation, thus we set the output component numbers to 15 and 30 and found that the 30 component analysis more clearly defined components whose loadings were related to SSWV/study. Hence results of the ICA denoising method for CT were based on the ICA with 30 components. Among those, 11 components for right hemisphere CT and 9 components for left hemisphere CT were significantly related with SSWV and study variables and were thus removed from the original data. With no denoising, there were many regions with significant group differences that were related to SSWV and study variables. GLM regression, ICA, and LICA-SR denoising methods remove some noise-related group differences. LICA-R2 shows better denoising performance with only a few regions with significant group differences remaining after LICA-R2 denoising, while LICA-R1 denoising results in the cleanest maps.

**Figure 6.**
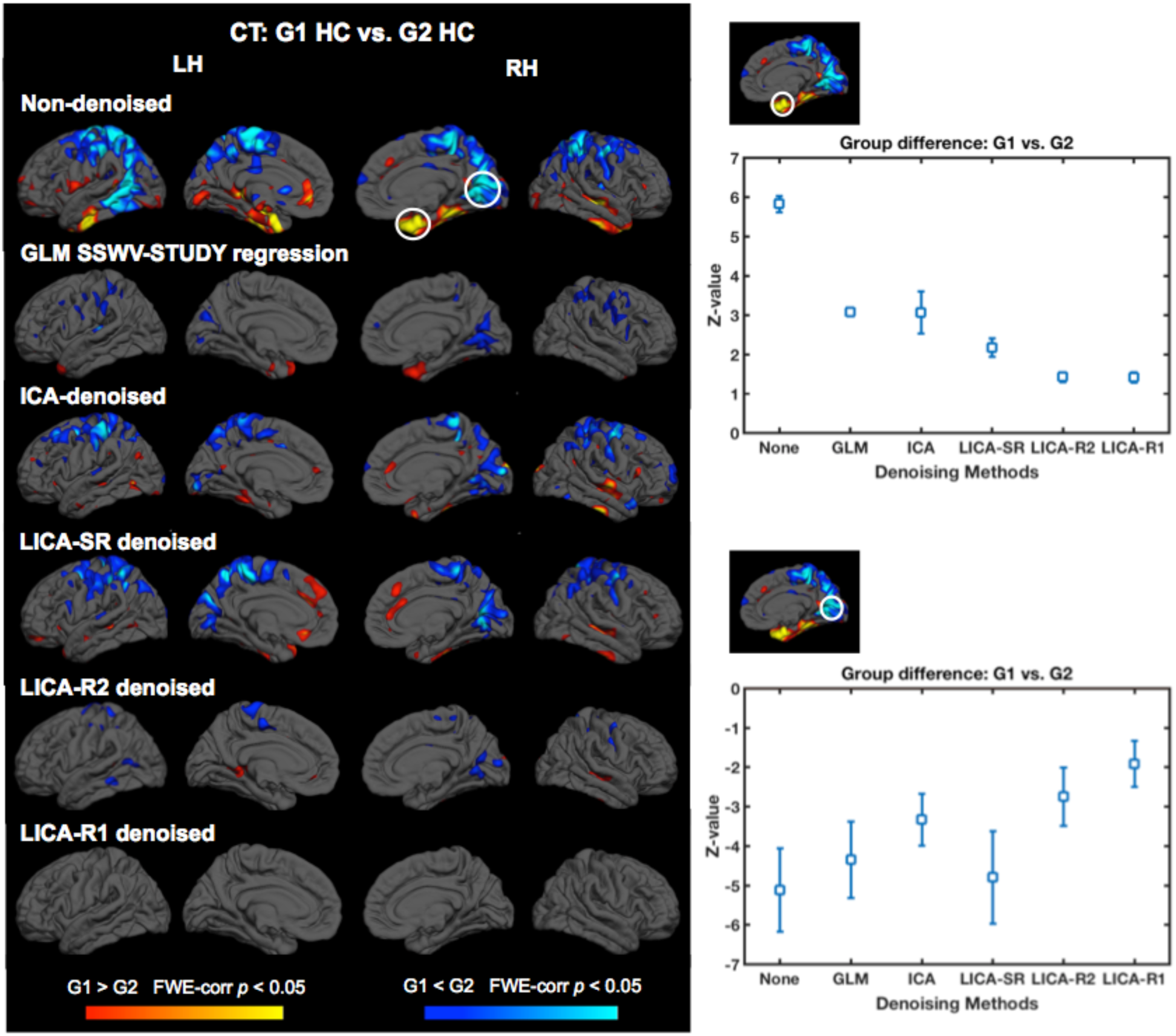
Group-level analysis of CT data before and after denoising. Two groups of HC data were constructed based on SSWV. G1 contains 29 subjects with data acquired pre-TIM upgrade; G2 contains 42 subjects with data acquired post-TIM upgrade. Resulting group difference maps without any data denoising (first row) and with data denoising (rows 2-6) and plots (right) of the mean Z-values and standard deviations extracted from voxels with significant group differences (shown in the white circles) show results that are similar to the GM and fMRI results in that every denoising procedure removes some of the effects of SSWV, while LICA-R1 denoising achieves the cleanest group difference maps.

Fig. 7 shows the correlation between subject loadings and subject-specific regression weights obtained from the first stage regression of the LICA noise spatial maps against each modality’s subject series conducted for LICA-R2 (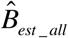 from Equation 2) for the three noise components from the LICA. Subject-specific regression weights from the first stage regression of the noise LICA component spatial maps against each the data for each individual modality (e.g., with a different set of regression weights for component and for each modality) are not equivalent to the component subject loadings, but there is a high correlation between them. For the first two noise components, which are multimodal, the correlation levels were very high for each modality. For the third noise component, the correlation is especially high for GM (with r-value nearly equal to 1), with less correlation between subject loadings and regression weights for the other modalities, which is due to the strong weighting toward GM for this pattern with very little contribution from other modalities.

**Figure 7.**
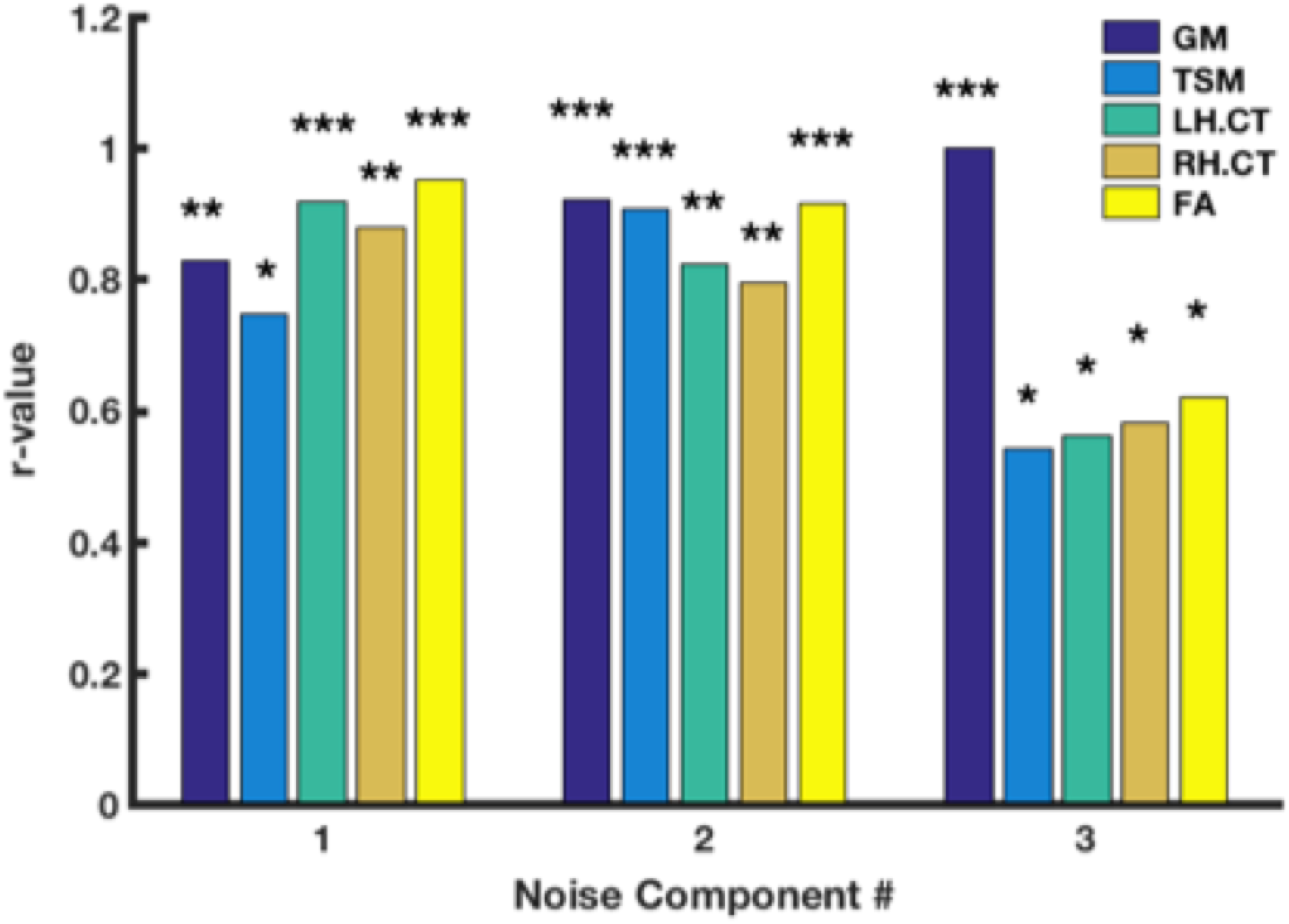
Correlations between LICA subject loadings for each noise component and the subject-specific regression weights for each modality obtained from the first stage regression of the LICA noise spatial maps against each modality’s subject series conducted for LICA-R2. Subject-specific regression weights are not equivalent to the LICA subject loadings, but they are highly correlated for modalities represented in any given component. For example, for multimodal noise components #1and #2, correlations were very high for all modalities. For noise component #3, which is primarily weighted on GM, the correlation is very high for GM, but reduced for other modalities. * *p* < 10^-5^, ** *p* < 10^-30^, *** *p* < 10^-50^.

## Discussion

A data denoising approach was developed to remove scanner and study variability from multimodal MRI data to facilitate combining multi-study data that is also applicable to combining multi-site data. Our approach utilizes a data fusion technique, LICA (Groves et al., 2011), to identify multimodal spatial components and a corresponding set of subject loadings for each component that are related to scanner/study effects. Because either spatial patterns or subject loadings can be used to denoise the data, we assessed three different strategies for removing the noise components identified by LICA: two “hard” or aggressive denoising methods - LICA-R2 (dual multivariate regression utilizing only the LICA noise spatial maps) and LICA-R1 (single multivariate regression utilizing only the LICA subject loadings for the noise components), and one “soft” regression-based denoising method - LICA-SR (single multivariate regression using all LICA component subject loadings). We also compared the performance of LICA-based denoising with two other approaches/analytical methods used when combining multi-site/study data; 1) GLM regression that included site/study covariates in the group-level model (SSC-GLM) to statistically control for scanner/study effects *post hoc* and 2) single-modality ICA denoising prior to group level statistical analysis. Overall, LICA-R1 performed well across all modalities and brain regions, demonstrating superior performance when compared with the other methods we tested by reducing SSWV and study-related variance while not introducing false positives.

LICA-based denoising is optimally suited for denoising multimodal MRI data for several reasons. The first is that noise-related effects are identified with a multivariate data-driven method that utilizes the shared covariance among the modalities (i.e., each subject has multiple measurements) to identify linked structure-function patterns and their corresponding subject loadings (measures of the degree to which a multimodal spatial pattern is present in a given subject) that capture both real “signal” and noise sources reflecting between-subject variability. The second is that removal of the noise effects identified by LICA proceeds with multivariate regression. This is in contrast to the SSC-GLM regression method that attempts to statistically control for confounds *post hoc* by including simple scanner/study confound regressors in the higher-level voxel-wise GLM. In this approach, the covariates do not capture any between-subject variability within site/study (for example, day-to-day variations in scanner performance at a given site or during a particular study), and it treats each voxel as being independent of all other voxels when controlling for such effects. Ignoring spatial covariance, as is done in voxel-wise GLM, is an unrealistic assumption for analyzing brain morphology or function that likely results in reduced sensitivity to detect effects when compared with multivariate methods that utilize spatial covariance information. While voxel-wise GLM is relatively easy to implement (but see Glover et al., 2012) and is modestly effective for some modalities, Glover et al. (2012) identified a number of weaknesses in this approach that are addressed using our proposed approach.

LICA denoising also offers advantages for multimodal MRI data as compared with modality-specific ICA-based denoising. Modality-specific ICA-based denoising identifies noise components based only on a single modality and thus do not capitalize on shared covariance among modalities to identify scanner/study effects. In addition, while LICA and single-modality ICA both require estimating the dimensionality (or number of components) and manually identifying noise signals, this only has to be done once for LICA, whereas for single-modality ICA, it has to be done for each modality (i.e., each subject-series) separately. Ideally both LICA and ICA dimensionalities should be “tuned” to give the best separation of signal and noise into individual components (that is, run at different dimensionalities to identify the best separation), a process that is relatively easier to do one time for LICA rather than doing several times for many individual modalities. It should also be noted that for fMRI data, it is common to denoise each subject’s fMRI timeseries data, which only removes within-subject sources of noise, such as motion and artifacts. While there may be scanner-related noise during that fMRI run, which could be identified and removed by ICA denoising, this would be an inefficient way of doing so as it would not capture between subject variability associated with software/hardware upgrades or multi-site effects. Thus, if using single-modality ICA to denoise scanner/study effects from fMRI, one should apply this method to the subject-series (activation maps for all subjects together), not the single-subject fMRI data itself. However, our findings show that this method is not as effective as multimodal denoising.

We also tested three different strategies for removing noise signals identified by LICA from the data for each modality: two aggressive denoising methods using the LICA subject loadings or the spatial maps for only the noise components (LICA-R1 and R2, respectively) and one nonaggressive method using the LICA subject loadings for all components (LICA-SR). We found that aggressive denoising using the subject loadings (LICA-R1) from LICA results performed consistently and generally better than other methods, including aggressive denoising using the LICA spatial maps and non-aggressive denoising using all LICA subject loadings. This is due to the fact that aggressive methods remove all the noise-related variance completely, and the subject loadings of the noise components capture within-scanner/study sources of variability in addition to the inter-scanner/study effects. Aggressive denoising using the LICA spatial maps appears to be slightly less effective than using the loadings (for removing noise effects), possibly because the subject-specific regression weights from the first stage regression (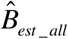 from Eq. 2) should be equivalent to the component subject loadings from LICA, but are not (because the regression is done for each modality separately whereas loadings are linked across modalities). The correlation between the LICA loadings and the subject-specific regression weights were extremely high when the noise component being removed was heavily weighted to a single modality that was the same as the modality being denoised, but was reduced when using these same components to denoise other modalities that did not contribute as much to the LICA noise pattern. Thus, using the LICA spatial maps to derive modality-specific regression weights appears to be as effective at removing noise as LICA-R1, but because the regression weights used in the second stage of LICA-R2 are now modality specific, LICA-R2 may not as aggressive as LICA-R1. In principle, soft regression-based denoising uses the subject loadings for all LICA components (LICA-SR), which removes only the unique variance related to the noise components to achieve a balance between noise removal and signal loss. However, we found that this approach is not as effective in removing scanner/study noise.

Although the data we used to test the LICA denoising approach were collected on one 3T scanner, the data have the same sources of variance in them that may be found in multi-site/multi-study data. For example, our data differ in the software versions, imaging acquisitions, and were collected via both pre- and post-TIM upgrade (which involved upgrading to a new gradient coil and from an 8-channel head coil to a 32-channel coil). Thus, scanner and study effects that are inherent confounds in multi-site data are similarly present in our multi-study data. The proposed LICA-based denoising methods are suitable for pooling data collected across different sites or across different operating systems on the same scanner, including field strength, scanner vendors, major scanner upgrade, and pulse sequence.

A limitation of this study is that although we tested the performance of different denoising methods by separating healthy control data into two groups based on SSWV or study variables that were matched for age and sex, other variables may have an effect on the differences between groups in these comparisons (observed using non-denoised data in Figs. 4-6). However, GM, fMRI, and CT values were significantly associated with SSWV or study variables and the regions with statistically significant differences with no denoising were not significant after LICA-based denoising of *only* scanner/study effects. Thus the effects associated with age, sex or other demographic factors are likely not contributing to the group differences.

## Conclusions

Scanner software and hardware upgrades and use of different acquisition parameters (e.g., study effects) can have a significant effect on the resulting MRI data, giving rise to the potential of misinterpreting findings derived by combining data from multiple sites or studies. Given that most studies collect multimodal MRI data over a period of many months or years, scanner software/hardware version changes can be a prevalent source of variability within a given study that can vary by modality and statistical outcome and reduce sensitivity to detect effects and/or induce false positive findings. We have developed and conducted initial testing of a new application of LICA for removing such effects from multimodal MRI data to overcome some of the limitations of existing methods for combining multi-site/study MRI data. Using a multivariate data-driven method for identifying sources of site/study-related variability, LICA, combined with a multivariate regression approach for denoising data, provides a strategy that more accurately and more efficiently identifies and removes site/study effects even when compared to using a higher level GLM with site/study covariates or single-modality ICA denoising techniques. This strategy will prove useful as the need for analyzing big data and the conduct of large multi-site projects continues to grow.

## Acknowledgements

This work was supported by the National Institutes of Health grant DA037265 (LDN). Data collection was supported by National Institutes of Health grants DA016695 (SG), DA021241 (SG), DA024007 (SEL), DA029115 (KPH) and AA014651 (MMS), and by DARPA-12-12-11-YFA11-FP-029 (WDSK).

